# A Real-time Multi-Subject Three Dimensional Pose Tracking System for Analyzing Social Behaviors of Non-human Primates

**DOI:** 10.1101/2024.02.27.582429

**Authors:** Chaoqun Cheng, Zijian Huang, Ruiming Zhang, Guozheng Huang, Han Wang, Likai Tang, Xiaoqin Wang

**Affiliations:** Tsinghua Laboratory of Brain and Intelligence, Tsinghua University, Beijing, China; Department of Biomedical Engineering, Johns Hopkins University, Baltimore, MD, US.

## Abstract

The ability to track positions and poses (body parts) of multiple monkeys in a 3D space in real time is highly desired by non-human primate (NHP) researchers in behavioral and systems neuroscience because it allows both analyzing social behaviors among multiple NHPs and performing close-loop experiments (e.g., delivering sensory or optogenetics stimulation during a particular behavior). While a number of animal pose tracking systems have been reported, nearly all published work lacks the real-time analysis capacity. Existing methods for tracking freely moving animals have been developed primarily for rodents which typically move on a 2D space. In contrast, NHPs roam in a 3D space and move at a much faster speed than rodents. We have designed a real-time 3D pose tracking system (MarmoPose) based on deep learning to capture and quantify social behaviors in natural environment of a highly social NHP species, the common marmosets (*Callithrix jacchus*) which has risen to be an important NHP model in neuroscience research in recent years. This system has minimum hardware requirement and can accurately track the 3D poses (16 body locations) of multiple marmosets freely roaming in their homecage. It employs a marmoset skeleton model to optimize the 3D poses and estimate invisible body locations. Furthermore, it achieves high inference speed and provides an online processing module for real-time closed-loop experimental control based on the 3D poses of marmosets. While this system is optimized for marmosets, it can also be adapted for other large animal species in a typical housing environment with minimal modifications.

## Introduction

The common marmoset (Callithrix jacchus) has emerged in recent years as a promising non-human primate model in neuroscience research, offering unique advantages over other animal models. Compared to rodents, marmosets have more complex brain architecture and exhibit closer cognitive ability to humans. Unlike larger primates like macaques, marmosets are easier to breed in captivity and have a shorter developmental stage and faster reproductive cycle(*1, 2*). Marmosets have been widely used in various fields of scientific research, including vocal and auditory studies(*3-8*), visual neuroscience(*9-11*), and transgenic studies for disease modelling(*12-16*).

Marmosets are particularly suitable for a wide range of behavioral experiments due to their small body size and social behaviors(*17*). However, most behavioral experiments on marmosets have been conducted based on manual recordings or with movement constraints(*18-20*). Therefore, the ability to automatically capture and quantify behaviors of marmosets in natural environment and social scenarios is highly desired by marmoset research community. Such a system could be integrated with other experimental methodologies to advance marmoset research. For instance, using quantified behaviors to identify the differences in behavioral phenotypes between normal and genetically modified marmosets could shed light on the relationships between genes and behaviors. In addition, synchronizing behavioral quantification with neural activity recording technologies have the potentials to reveal the neural mechanisms underlying behaviors.

In recent years, there has been a rapid development of automated pose tracking systems for animal behavioral studies. DeepLabCut offers multi-animal 2D pose estimation capabilities(*21, 22*), and can be extended to provide low-latency 3D pose estimation for a single animal(*23-25*). SLEAP provides versatile support for multi-animal 2D pose tracking across a variety of network architectures(*26, 27*). DANNCE enables direct 3D pose estimation for single rodent and can be transferred to other species(*28*). MAMMAL provides the capability to capture 3D surface motions of pigs and dogs(*29*). Additionally, species-specific systems have been developed, such as OpenMonkeyStudio for macaques(*30*), DeepFly3D for drosophila(*31*) and FreiPose for rats(*32*).

DeepBhvTracking(*33*), MarmoDector(*34*) and FulMAI(*35*) are offline tracking systems specifically designed for marmosets. However, the functions of these systems are confined to tracking the positional trajectories of marmosets in 2D or 3D spaces, lacking the crucial pose information for behavioral analyses. While DeepLabCut offers 2D pose tracking capability for marmosets, and DANNCE has the potential to be adapted for estimating 3D poses of single marmoset, a significant amount of new training data are required for new experimental setups, such as adding more marmosets with new identities, which considerably limits their applications in marmoset behavioral experiments.

Due to the highly social nature and rapid movements of marmosets in three-dimensional space, a system capable of tracking 3D poses of multiple marmosets is highly desired. In addition, the ability to control experimental stimuli in real-time based on the marmoset’s positions or actions(*25*) would give researchers power to conduct a wider range of behavioral and physiological experiments in freely roaming marmosets. Existing systems are unable to fulfill these specific requirements. In this study, we have developed an efficient and user-friendly real-time 3D pose tracking system for multiple marmosets, which can be flexibly adapted by a wide range of researchers to study the marmoset’s natural behaviors.

The MarmoPose described in this report is a deep learning-based 3D pose tracking system, with minimum hardware requirement, specifically designed for reconstructing 3D poses of single or multiple marmosets freely moving in their homecage environment. In MarmoPose, multi-view images captured by four (or more) cameras are first processed by deep neural networks to predict 2D coordinates of 16 body locations of each marmoset. Subsequently, visible 3D body locations are reconstructed using triangulation while invisible 3D body locations are estimated through a denoising autoencoder incorporating a marmoset skeleton model. MarmoPose offers several advantages over existing systems: (1) This is the first system to enable comprehensive 3D pose tracking for multiple marmosets; (2) This system supports real-time closed-loop experimental control based on the 3D poses and positions of marmosets, which could be integrated with other experimental functions including stimulus playback and neural recording; (3) This system employs a marmoset skeleton model for 3D coordinates optimization, thereby improving the precision of the reconstructed 3D poses and rendering it possible to estimate invisible body locations in the cameras’ blind spots; (4) This system is flexible as each module can be independently modified to accommodate new experimental setups, such as varying the number of marmosets in the cage or different obstacle configurations; (5) This system is designed for user-friendly deployment in a typical marmoset family cage (0.7m x 1m x 0.8m) without additional modifications and can therefore be easily adapted to other housing or experimental environments.

## Results

### Overview of the MarmoPose system

MarmoPose is a 3D pose tracking system specifically designed for both single and multiple marmosets. It receives video streams from multiple cameras and outputs estimated body locations of marmosets in a 3D space. We developed the system for a typical marmoset family cage (1m x 0.7m x 0.8m) with four video cameras mounted on the upper corners as shown in Fig. 1a. Marmosets can freely move around in the cage. Two wooden logs, two mental shelves and some small sticks are placed inside the cage to enrich the environment. Note that the cameras are fixed on four top corners inside the homecage so that such a setup does not require any modifications of the cage and thus can be easily deployed in other housing cages.

**Fig. 1.**
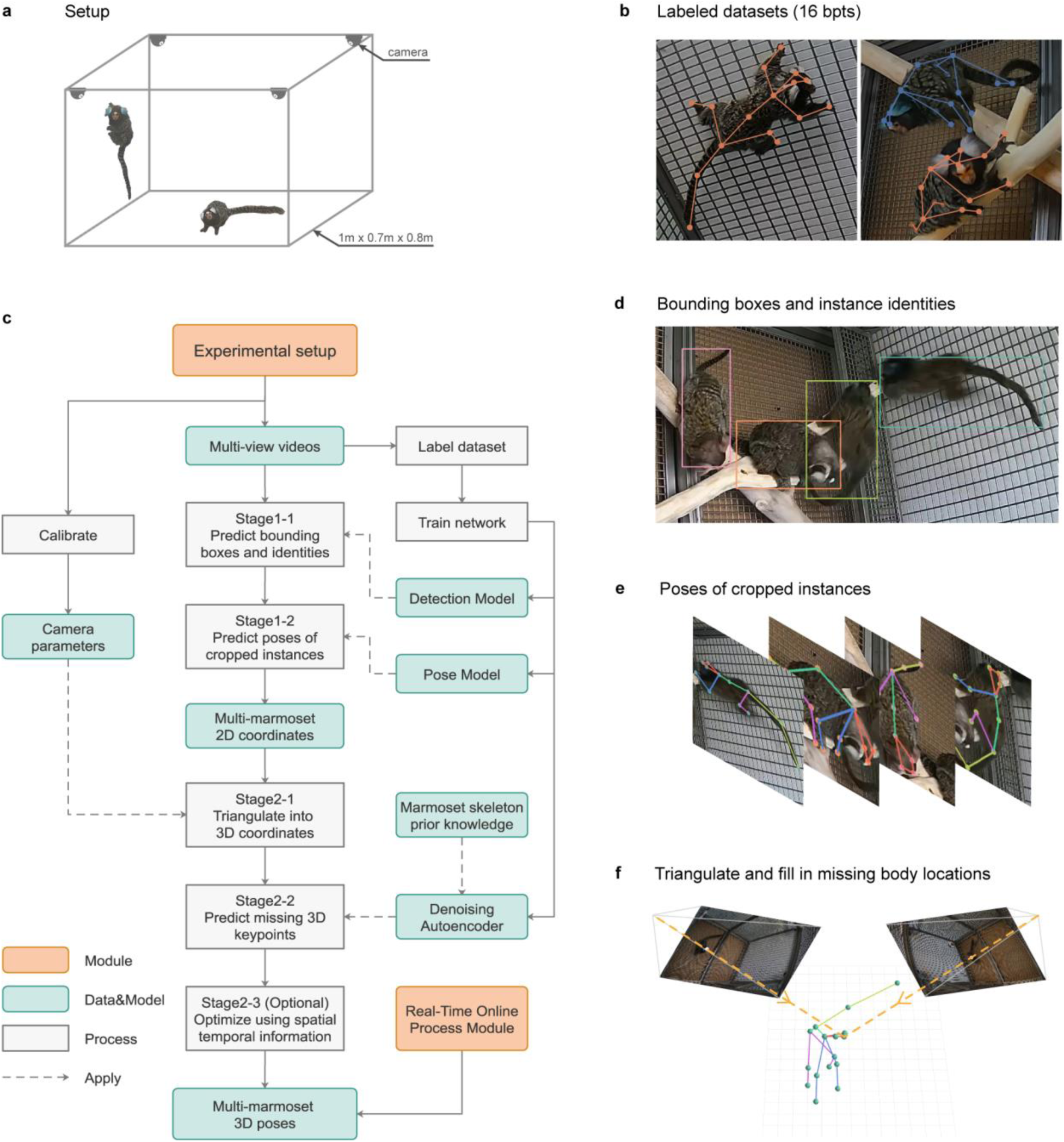
MarmoPose is a complete system for multi-marmoset 3D pose tracking. **a**, Experimental setup of MarmoPose. Four cameras are mounted on the upper corners of the marmoset homecage (1m x 0.7m x 0.8m), and two wooden logs and two mental shelves are placed in the cage as their daily environment. Marmosets can freely move without any interference. **b**, Diagram of the whole workflow of MarmoPose. **c**, Example (cropped) images with manual annotations for the Marmoset3K dataset. Each dot represents one of the 16 body locations (transparent dots denote invisible body locations from this camera view), and the lines indicate the marmoset skeleton. **c**, Diagram of the whole workflow of MarmoPose. **d**, Example (cropped) image with predicted bounding boxes, where the instance identity of the marmosets is denoted by the color of the bounding box. **e**, Example (cropped) images with predicted body locations, where invisible body locations are omitted. **f**, Illustration of 3D triangulation with 2D predictions.

We selected 16 locations of the body to capture the posture of a marmoset (head, left/right ear, neck, spinemid, left/right elbow/hand/knee/foot, tailbase, tailmid, tailend). As annotated in Fig. 1b, each dot represents one of the body locations and the lines represent the marmoset skeleton. Invisible body locations from this view are indicated by transparent dots. In scenarios involving multiple marmosets, maintaining consistent identification of each individual across different camera views is crucial for accurate 3D triangulation. To ensure reliable identification of individuals, we applied a harmless dye to their ears to clearly distinguish them (For *n* marmosets, *n−*1 are marked by different colors while one is unmarked). For instance, the marmoset marked with blue dye is correspondingly annotated with blue in Fig. 1b.

Fig. 1c illustrates the workflow of MarmoPose, consisting of two main stages. In the first stage, 2D predictions were generated for each video. For images containing multiple marmosets, we first trained a detection model, adapted from RTMDet(*36*), to detect the bounding box as well as the identity of each marmoset (Fig. 1d). Subsequently, we trained a pose estimation model, adapted from RTMPose(*37*), to predict 16 body locations based on the images cropped around these bounding boxes (Fig. 1e). We adopted the two-stage approach due to its flexibility in accommodating new experimental setups and the relatively small size of marmosets in each camera view. To train these deep neural networks, we labeled the Marmoset3K dataset (see Methods) based on 1527 images containing one marmoset and 1646 images containing two marmosets across different camera views. Each image was annotated with the 16 body locations and identity information for each instance. In the second stage, instances with the same identity from all camera views were integrated and triangulated into 3D poses using camera parameters (Fig. 1f) which were calibrated once the cameras were fixed (see Methods). Subsequently, we employed a denoising autoencoder (DAE)(*38*) integrated with prior knowledge of a marmoset skeleton model to reconstruct invisible body locations (see Fig, 3c). Finally, an optional step could be performed to refine the 3D poses further using an iterative optimization method (see Methods). As a result, MarmoPose can accurately estimate 3D poses of the marmosets with both visible and invisible body locations. To help visualize the estimated 3D poses, MarmoPose also generates videos combining images from camera views with predicted body locations and identity information (Fig. 2a). In addition, MarmoPose provides an online process module to enable real-time experimental control based on the 3D poses of marmosets. This module will be discussed in detail below.

**Fig. 2.**
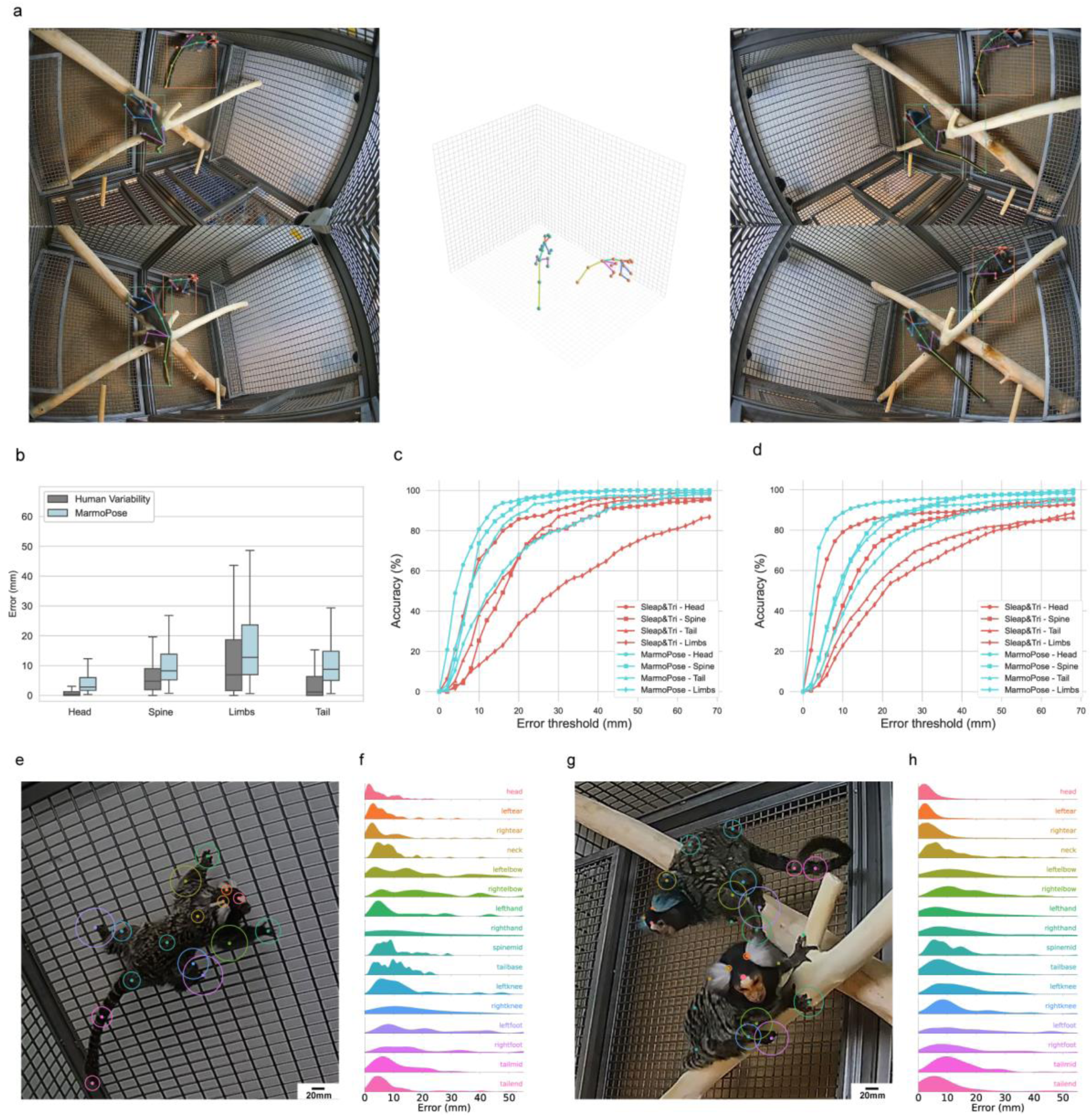
MarmoPose can accurately track 3D poses of multiple marmosets. **a**, An example demonstrating one frame of the visualized videos generated by MarmoPose. Four corners show images from 4 camera views with predicted 2D body parts and identities, while the central part shows the reconstructed 3D poses. **b**, Box plots of 3D Euclidean errors for MarmoPose and human variability evaluated on the Marmoset3D test dataset, 16 body locations are grouped into 4 categories based on their anatomical locations (Head: head, left/right ear; Spine: neck, spinemid; Limbs: left/right elbow/hand/knee/foot; Tail: tailbase, tailmid, tailend). Human variability indicates the error between hand-labeled ground-truth on the same data from two different people. (n=258 instances). **c**, 3D reconstruction accuracy as a function of error threshold evaluated on the test data (n=71 instances) for single marmoset. Body locations are broken down to the same categories as in **b**. **d**, 3D reconstruction accuracy as a function of error threshold evaluated on the test data (n=152 instances) for paired marmosets **e-h**, 3D reconstruction errors for single marmoset and paired marmosets. Each dot represents one of the body locations, with surrounding circles denoting the 75th percentile of the 3D Euclidean errors, and histograms correspond to full error distribution evaluated on the test data (n=84 instances for single marmoset dataset; n=146 instances for paired marmosets dataset)

### MarmoPose can accurately track 3D poses of multiple marmosets

Fig. 2a shows a visualized frame generated by MarmoPose. Original images from the four cameras are annotated with predicted identities, bounding boxes and 2D poses of two marmosets in the cage. The central plot shows the reconstructed 3D poses of the two marmosets, reflecting the marmosets’ 16 body positions and poses as they roam freely in the cage. This figure demonstrates that MarmoPose can track the 3D poses of multiple marmosets (see Movie 1).

In order to evaluate the accuracy of MarmoPose quantitatively, we constructed a dataset named Marmoset3D (see Methods) by triangulating hand-labeled 2D coordinates from multiple camera views at the same timepoint. The Marmoset3D dataset contained 522 3D ground-truth instances with 8352 body locations (16 body locations per each instance), consisting of 140 instances collected from 140 timepoints with single marmoset (from four marmosets) and 382 instances collected from 191 timepoints with paired marmosets (from three pairs of marmosets). The hand-labeled 2D coordinates were first annotated by one person and then proofread by a second person to ensure accuracy. However, because marmosets are relatively small in each camera view and there were no obvious landmarks on their bodies for precise localization, there was inevitably variability in the 2D coordinates annotated by the two persons. Theoretically, this human variability is the lower bound of the error of MarmoPose. We computed the human variability by measuring the error between hand-labeled ground-truth on the same data from two different persons. Fig. 2b shows the 3D Euclidean errors of MarmoPose and human variability, in which 16 body locations are grouped into 4 categories based on their anatomical locations for clarity (Head: head, left/right ear; Spine: neck, spinemid; Limbs: left/right elbow/hand/knee/foot; Tail: tailbase, tailmid, tailend). MarmoPose can achieve comparable 3D errors to the human variability, the median error of human variability for these 4 groups are: 0.29mm(Head), 4.77mm(Spine), 6.92mm (Limbs) and 1.12mm(Tail), and the median error of MarmoPose for these 4 groups are: 2.82mm(Head), 8.25mm(Spine), 12.76mm(Limbs) and 8.75mm(Tail).

Given MarmoPose is the first complete system to track 3D poses of multiple marmosets, for comparison, we trained models using SLEAP with the same dataset to estimate the 2D poses of multiple marmosets, followed by triangulation using direct linear transformation from four camera views. Fig. 2c and Fig. 2d illustrate the 3D reconstruction accuracy as a function of error threshold for both MarmoPose and SLEAP combined with triangulation (abbreviated as Sleap&Tri) evaluated on the Marmoset3D test dataset of single marmoset and paired marmosets respectively, where the accuracy is defined as the percentage of 3D body locations with 3D Euclidean errors below the error threshold, with 16 body locations grouped into four categories. The figures clearly demonstrate that MarmoPose outperforms Sleap&Tri significantly. Quantitatively, the typical body length of an adult marmoset is about 20cm (excluding the tail, which is also approximately 20cm long). If we choose 20 mm (about 10% of body length) as a threshold, the accuracy of MarmoPose is 95% for Head, 93% for Spine, 68% for Limbs and 89% for Tail respectively on the single marmoset test data (Fig. 2c), and the accuracy of MarmoPose is 94% for Head, 85% for Spine, 68% for Limbs and 83% for Tail respectively on the paired marmosets test data (Fig. 2d). Note that even with the largest threshold analyzed, the accuracy still can’t reach 100%. This is partly due to occasional incorrect identification of marmoset identities, leading to significant errors after 3D triangulation.

Figure 2e and 2g display the 3D reconstruction errors with two representative images. The 75th percentile of the 3D Euclidean errors for each body location is represented by a circle The error distributions are shown in Fig. 2f and 2h. Generally, body locations on the head have smaller errors due to the existence of boundary between brown and white hairs, while body locations on the limbs have larger errors due to high degrees of freedom and frequent occlusion.

### MarmoPose can be easily adapted to new experimental setups

MarmoPose provides a detection model and a pose estimation model pre-trained on the Marmoset3K dataset, containing images of single and paired marmosets. When conducting experiments, researchers may need to deploy MarmoPose in new experimental setups different from the default configuration, for example, with a different number of marmosets and additional items inside the cage. As demonstrated in Fig. 3a, MarmoPose can be easily adjusted to accommodate these changes by finetuning one of the models with minimal additional effort. For instance, we conducted a test to track a family of four marmosets with new identities (marked by new dye colors). Under this circumstance, only the detection model requires finetuning, and only the bounding boxes with corresponding identities for each instance need to be labeled in the new training data. In fact, we annotated 100 images (which took less than 1 hour) to finetune the detection model. As a result, the finetuned model can achieve 93.2% identification accuracy with precise bounding boxes (Fig. 3b), which can be directly used in experiments involving four marmosets. Another issue is that the dye colors marked on marmosets will inevitably be occluded by obstacles or become invisible in some camera views. To mitigate this issue, we designed a heuristic assignment algorithm (see Methods) to assign the identity labels predicted by the detection model for each instance, which can further increase the accuracy from 95.1% to 96.5% for two marmosets and 93.2% to 95.0% for four marmosets, as depicted in Fig. 3b.

**Fig. 3.**
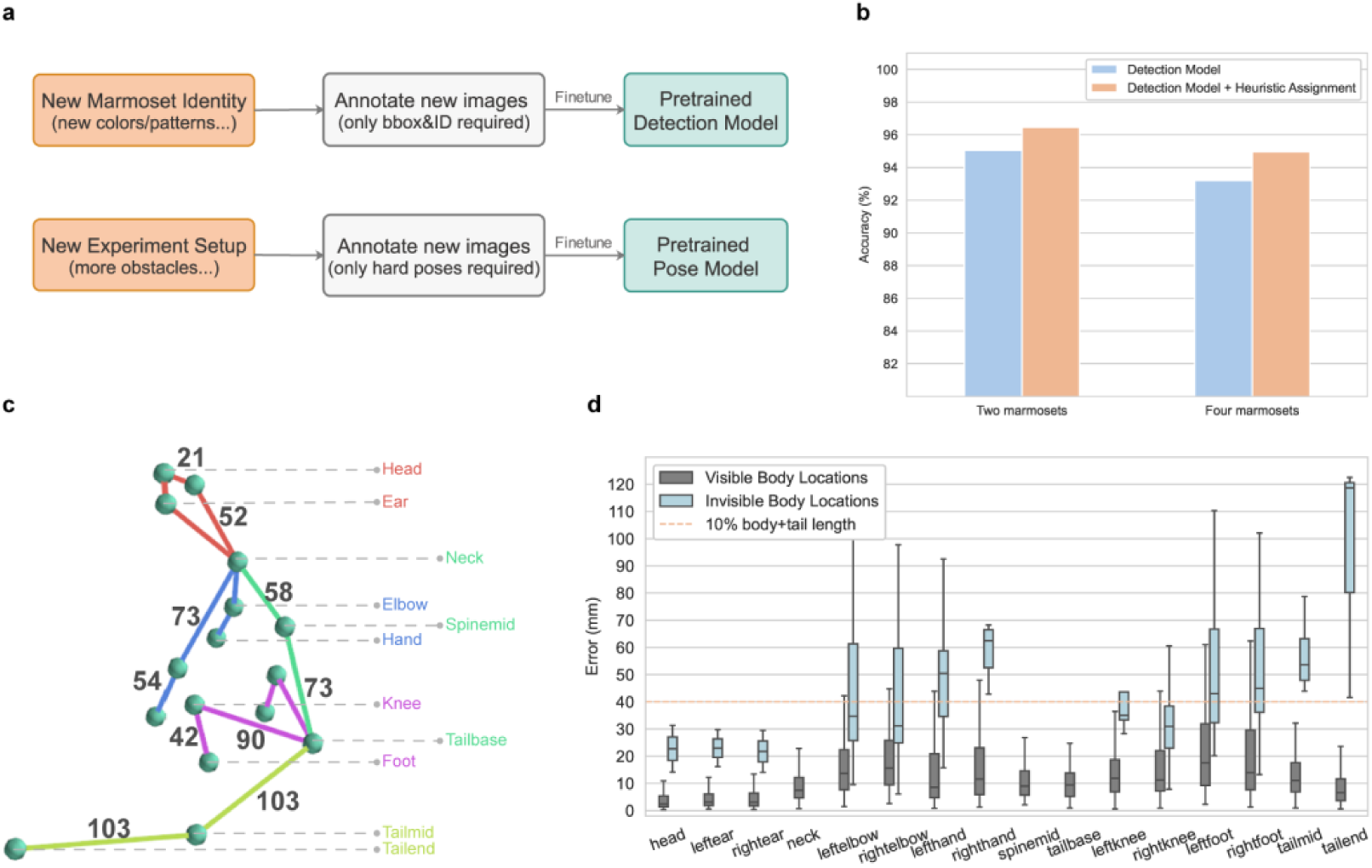
MarmoPose can be easily adapted to new experimental setups and can estimate invisible body locations using denoising autoencoder. **a**, Schematic for transferring MarmoPose to new experimental setups through a two-stage workflow. For incorporating new marmoset identities (like new colors or other patterns), only bounding boxes and corresponding labels need to be annotated to finetune the pretrained detection model. For deploying in a complete new experimental setup (like a scenario with more obstacles), only poses in difficult situations need to be annotated to finetune the pretrained pose model. **b**, Identification accuracy of the detection model with and without heuristic assignment evaluated on paired marmosets dataset. **c**, Illustration of the marmoset skeleton model, used as one of the terms in the loss function of DAE to guide the reconstruction of missing body locations. Numbers indicating the median length of the distances between two body locations measured on real marmosets, with different weights assigned based on their degrees of freedom. **d**, Box plots of 3D Euclidean errors in test data for visible and invisible body locations. Note that ‘neck’, ‘spinemid’ and ‘tailbase’ have no invisible body locations because they are used to normalize the poses.

### MarmoPose can estimate invisible body locations using denoising autoencoder

In the scenarios involving multiple marmosets freely moving in the homecage, some of their body locations will inevitably be self-occluded or occluded by other animals and objects like logs or shelves in the cage, leading to missing data in the reconstructed 3D poses. Inspired by previous study on 2D human pose estimation(*39*), we trained a denoising autoencoder (DAE) to reconstruct the missing 3D body locations, which receives 3D coordinates of 16 body locations with missing data as input and outputs the estimated complete coordinates. We trained the model by randomly masking some body locations to simulate inputs with missing data based on the Marmoset3D dataset (see Methods).

In order to guide the denoising autoencoder to estimate missing coordinates better, we added an extra loss term to constrain the lengths between two body locations in the reconstructed 3D poses based on a prior knowledge of marmosets. We measured the median length of distances between two body locations on three normal adult marmosets to form a skeleton model (Fig. 3c). Incorporating this prior knowledge of marmosets into the model enhances its ability to capture the underlying structure of marmosets. As a result, the denoising autoencoder can more accurately reconstruct the missing body locations, even with a limited amount of training data. Fig. 3d shows the 3D error distributions for each body locations when they are visible (computed by triangulation of multiple 2D predictions) or invisible (estimated by DAE) evaluated on the Marmoset3D test data. Note that ‘neck’, ‘spindmid’ and ‘tailbase’ don’t have corresponding invisible values, because they are used to normalize the poses during the application of the denoising autoencoder. It can be observed that for the body locations on the head or spine, the errors of estimated coordinates are below 10% of the body and tail length (40mm), whereas the errors of tail positions are highest. This is reasonable because the marmosets’ tail is long and flexible, making their precise locations more difficult to predict. Although missing data predicted by DAE might not achieve the accuracy level of visible body locations, it still offers a reasonable estimation of the missing body locations within a controlled experimental setting.

### MarmoPose enables real-time experimental control based on 3D poses

One objective of the MarmoPose is to provide an online process module to enable real-time experimental control by processing images from multiple live video streams and output 3D poses frame by frame with a small latency. To achieve this goal, we adopted multi-processing and multi-threading to deal with different tasks in parallel. As the workflow chart shown in Fig. 4a, in one process, live video streams are first cached by multiple threads, then 2D detection and 3D reconstruction are performed on the latest cached frame. In the other process, poses and images are combined together for display and an interface returning the latest poses is provided for customized experimental control.

**Fig. 4.**
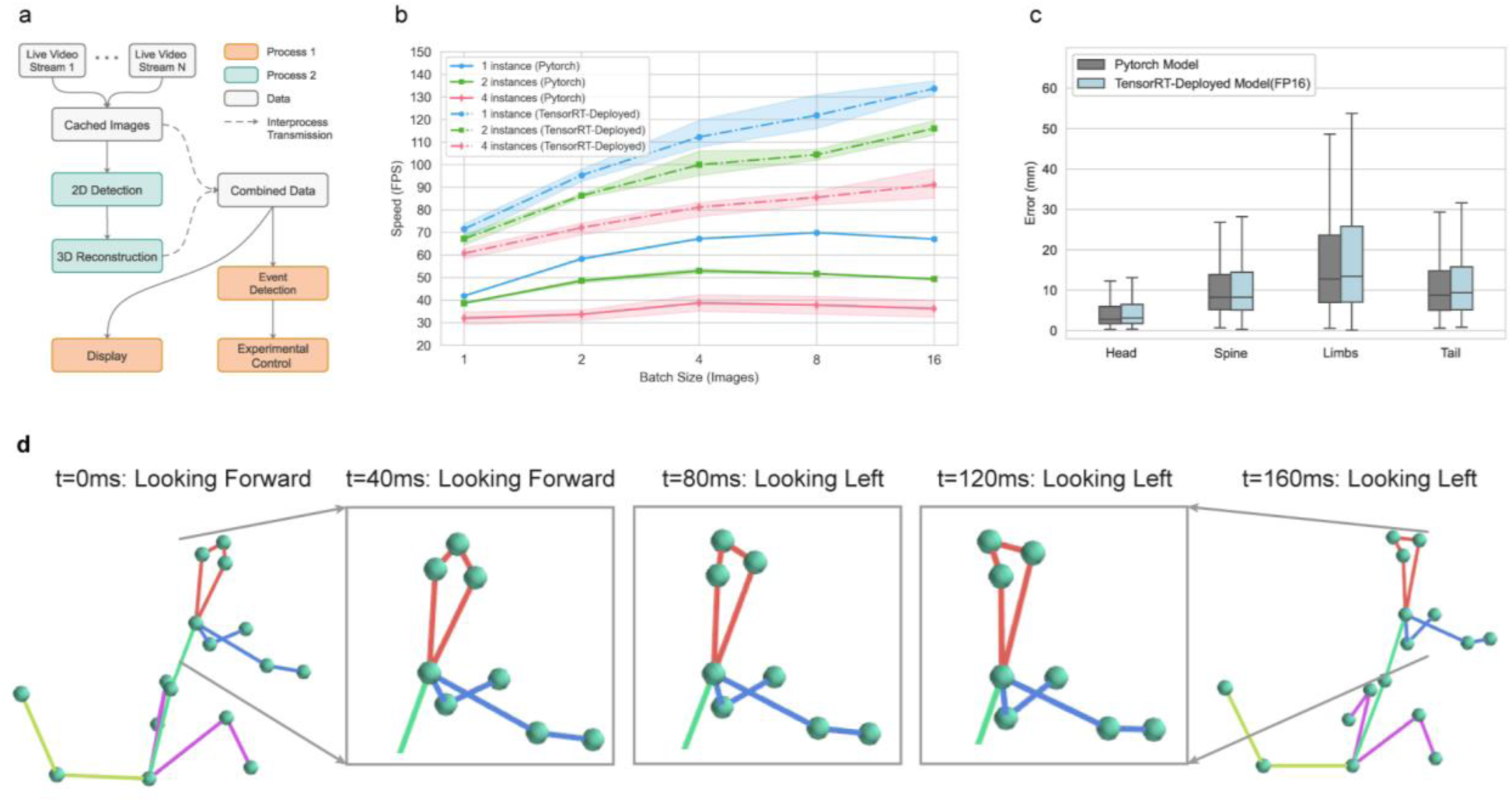
MarmoPose provides an online process module to enable real-time experimental control. **a**, Schematic of real-time experimental control. Process 2 reads images from cached live video streams and perform 2D detection and 3D triangulation, while process 1 (main process) reads images and poses to display and perform experimental control. **b**, Inference FPS variation across different number of instances in videos, evaluated over varying batch sizes, where each condition is evaluated on four different videos containing 1000 frames. Solid lines represent the original PyTorch model, while dashed lines denote the model deployed using TensorRT in FP16 mode. **c**, Box plots illustrating the 3D Euclidean errors of both the PyTorch model and the TensorRT-deployed model evaluated on the Marmoset3D test dataset. 16 body locations are grouped into 4 categories based on their anatomical locations. **d**, An example showing real-time event detection. The marmoset turns its head from forward to left in 160ms (4 frames), while MarmoPose could feedback the head orientation in less than 40ms and perform real-time experimental control based on the detected event.

The system latency of real-time processing module mainly consists of three parts: 2D pose estimation, 3D reconstruction and inherent processes such as data transfer and display, where 2D detection is the most time-consuming part. Actually, deploying the trained PyTorch model using TensorRT could significantly improve the inference speed on specific hardware. To compare the performance of the deployed models against the original PyTorch models. We conducted benchmarks on the combined inference speed of detection and pose estimation models. The benchmarks were performed across various number of marmosets appearing in the videos and different inference batch sizes. As illustrated in Fig. 4b, with a batch size of 4, the original PyTorch model achieves 68fps for videos with 1 instance, 53 fps for 2 instances and 39 fps for 4 instances. After the deployment via TensorRT, the inference speed was significantly increased to 112 fps for videos with 1 instance, 100 fps for 2 instances and 82 fps for 4 instances. For processing videos in real-time, we would combine the frames from different cameras into a pseudo batch for inference (see Methods). Consequently, the deployed model can seamlessly achieve real-time process by finishing all computation within 40ms (the time interval between two frames) in a typical setup involving four cameras and 2 marmosets. It is important to note that deploying the model in FP16 mode inevitably leads to some loss of accuracy. We evaluated the 3D distances errors of the original model and the deployed model across 4 groups of body locations on the Marmoset3D test dataset. The average accuracy loss (less than 7%) is reasonable given the significant speed improvement (Fig. 4c).

We tested MarmoPose in a scenario in which we want to detect the head orientation of a marmoset and provide real-time feedback to control the stimulus. Poses of the marmoset at different timepoints are shown in Fig. 4d. The marmoset looks forward at 40ms and then turns to the left at 80ms. With a minimum latency of less than 40ms here, MarmoPose can reliably track such quick movements in real-time, thereby providing possibilities for real-time experimental control.

### MarmoPose enables quantitative analysis of behaviors for multiple marmosets

A significant aspect of neuroscience research involves quantifying animal’s natural behaviors(*22, 28, 30, 40*). MarmoPose can be used to quantitatively analyze behaviors of both single and multiple marmosets in various social scenarios in a homecage environment. To demonstrate this capacity, we analyzed a 32-minutes video clip containing 48,000 poses of a single marmoset. We generated behavioral density maps and isolated clusters by applying the watershed transform(*41*) over a density representation of the t-SNE space (see Methods), where each cluster represents a typical behavior of the marmosets, including climbing, jumping and walking (Fig. 5a). This behavior map can be utilized to characterize the differences in behavior between different instances, or for a single instance across time or in different states.

**Fig. 5.**
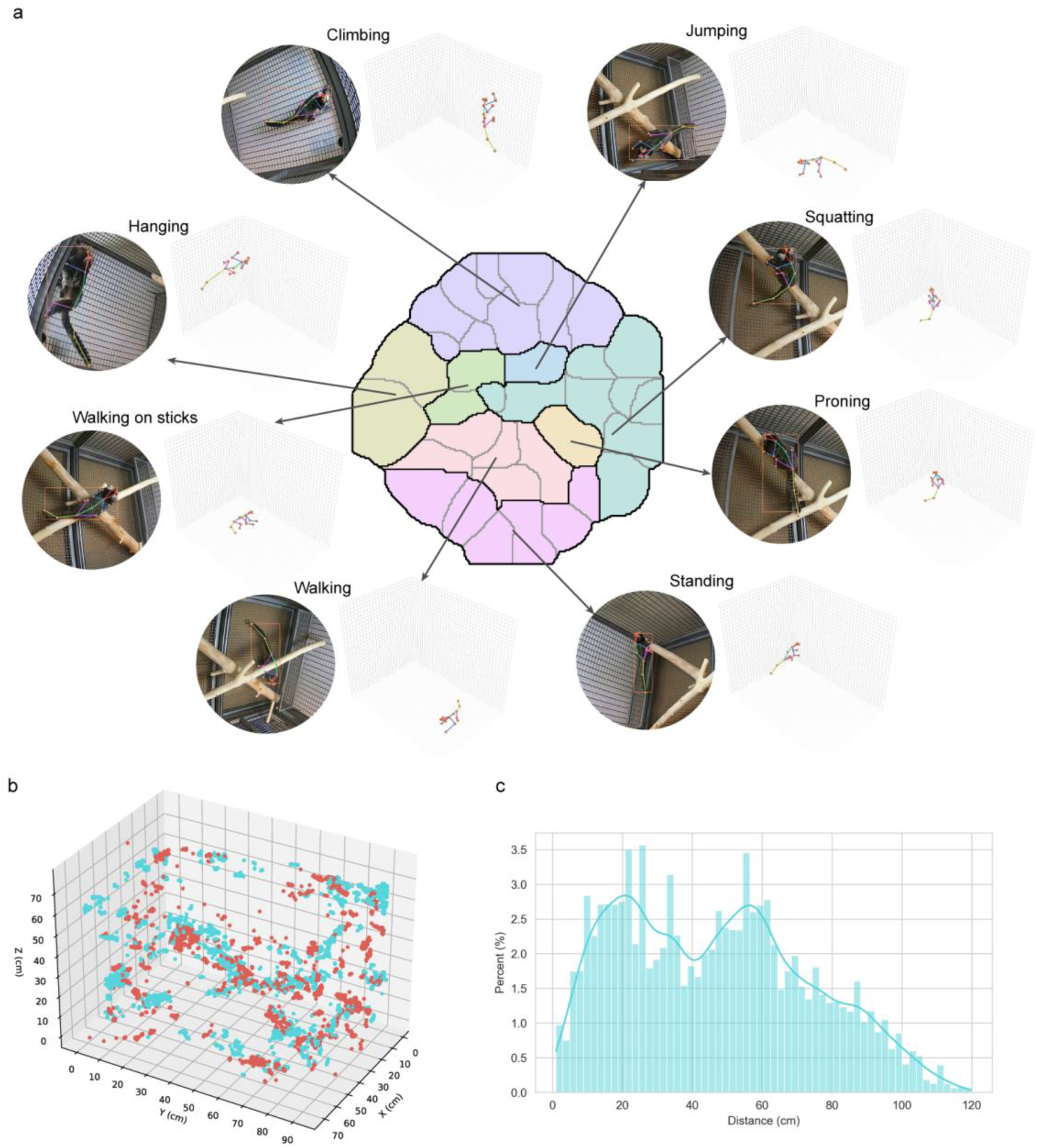
MarmoPose enables quantitative analysis of behaviors for multiple marmosets. **a**, The density map of t-SNE space overlapped with isolated posture blocks obtained from the watershed transform of the density map, with illustration of 8 distinct actions, including climbing, jumping, squatting, proning, standing, walking, walking on sticks and hanging. For each case, the left image is one of the raw images (cropped), and the right image is the reconstructed 3D pose. **b**, Color map of the duration each marmoset spent in different areas in the cage in 27 minutes. **c.** Distance distribution between two marmosets across 27 minutes.

Furthermore, we analyzed a 27-minutes video clip with two marmosets that were transferred into a new homecage environment. Fig. 5b illustrates the duration each marmoset spent in different areas of the cage. Notably, both marmosets frequently climbed the left and right walls, walked along the wooden logs, and rested on the two metal shelves, but they rarely stayed underneath these objects. Fig. 5c shows the distances distribution between the two marmosets during this period. The distance between these two animals peaked at ∼20cm and ∼60cm, respectively, indicating that the two marmosets didn’t engage in close social behaviors during this time while they explored the new environment.

## Discussion

Here we described MarmoPose that enables real-time 3D pose tracking for multiple freely moving marmosets in a housing cage environment. The primary considerations in designing the MarmoPose are to make it an efficient, cost effective (using minimum hardware), real-time and easily deployable in typical housing cages. This system is specifically designed for marmosets in order to optimize its performance. It has several distinct features comparing to other published pose detection systems for freely moving animals. First, MarmoPose leverages the prior knowledge of a marmoset skeleton model to estimate invisible body locations and refine 3D poses. Second, MarmoPose is a user-friendly system that can be deployed in a typical marmoset homecage environment and be easily adapted to new experimental setups. Third, MarmoPose provides a online process module for users to perform real-time closed-loop experimental control based on the 3D poses of marmosets, which can be integrated with other experimental methodologies such as stimulus playback and neural recording.

In scenarios involving multiple marmosets, particularly when they are in close proximity, distinguishing individual body locations accurately is a challenging task. This is a common problem in the field of multi-object pose tracking, which has not been addressed very well. Since each marmoset occupies only a small part of the whole image, and their rapid movements often lead to motion blur, it is challenging even for human annotators to accurately locate specific body locations. Consequently, MarmoPose might not perform optimally in such scenarios. A practical solution is to keep a balance between specific experimental demands and system accuracy. In many experimental situations, the focus might be only on a subset of body locations. Therefore, we can choose to ignore the misidentified data and concentrate on the relevant body locations. Additionally, in social scenarios, the overall interaction of the marmosets might be more significant than the precise poses of each body locations. Therefore, it might be more effective to treat marmosets that are close to each other as a single entity without attempting to distinguish every body part clearly.

In the current setup, four cameras are fixed on the upper corners inside the homecage to maximize coverage and minimizes maintenance and potential damages by animals. This is a simple and effective arrangement, which can be easily deployed in other housings without extra modifications, and keep the minimum computational cost at the same time. While adding more cameras to cover more blind spots is a straightforward solution to reduce invisible body locations and improve the system accuracy, it increases computing loads and slows down the processing speed. Practically, the number of cameras used should be determined by the specific experimental demands, the fewer the better. The current arrangement with four cameras is sufficient for a range of research purposes. For researchers aiming for higher precision, more cameras can be added to cover more blind spots, which is supported by MarmoPose. However, it is important to consider that increasing the number of cameras would lead to longer processing time and output latency.

The ability to automatically track poses of freely moving marmosets in 3D space in real-time using MarmoPose could significantly improve studies of natural behaviors in this field. Traditionally, studies on vocal communications(*4, 42*) and visual directional preference(*9-11*) have been relying on manually processed audio and video recordings. MarmoPose could be used to boost these studies by providing more accurate and comprehensive behavioral quantifications. Given their social nature and ease of handling, marmosets are ideal for comparative behavioral experiments with humans. For example, MarmoPose could be used to describe and compare the behavioral evolution and natural preferences between marmosets and human(*1, 17, 43*). Moreover, due to their relatively high reproductive cycles compared to other primates, marmosets are well-suited for transgenic modifications(*14, 44*) and disease modeling(*45-48*). Integrating MarmoPose with other methodologies such as neural recording and optogenetics offers possibilities to explore neural mechanisms underlying natural behaviors in both normal and genetically modified marmosets.

## Methods

### Datasets

We collected videos for single marmoset and paired marmosets to develop and evaluate MarmoPose. Videos were all recorded by four synchronized cameras (HIKVISION DS-2CD252CZY-ZFR) mounted on the upper corners of a typical marmoset homecage (1m x 0.7m x 0.7m) with 1920 *×* 1080 *×* 3 frame size at 25 fps. Two wooden logs, two mental shelves and some small sticks were placed in the homecage to encourage the marmosets to engage in naturalistic behaviors. We used the annotation tools provided by SLEAP(*27*) to initially label the ground truth, including 2D locations of 16 body locations for each marmoset and their identities. Subsequently, a python script was wrote to convert these labels into the COCO(*49*) format in order to train models incorporated within OpenMMLab.

All of the marmosets are healthy adults with normal body size and weight, and they have been lived in the same homecage for more than half a year. All of the procedures in these experiments were approved by the Science and Technology Ethics Committee at Tsinghua University.

### Marmoset3K - Single marmoset

Two human annotators labeled 1527 frames (1527 instances) from 4 different camera views across 16 videos capturing a single marmoset freely moving in the homecage. Four adult marmosets without additional modifications were used in this dataset. For each marmoset instance, 16 body locations were labeled (head, leftear, rightear, neck, spinemid, leftelbow, lefthand, rightelbow, righthand, leftknee, leftfoot, rightknee, rightfoot, tail-base, tailmid, tailend) carefully. If a body part is occluded from a camera view, it is labeled as invisible and will not be included in the training and testing process. To train the 2D pose predictor, we randomly selected 1270 frames as training data and the other 257 frames as test data.

### Marmoset3K - Paired marmosets

Three human annotators labeled 1646 frames (3292 instances) from 4 different camera views across 12 videos capturing a pair of marmosets freely moving in the homecage. Six (three pairs) adult marmosets were used in this dataset. Since the consistence of marmosets’ identities across different camera views is essential for accurate 3D triangulation, we dyed the ears of one marmoset in each pair with harmless blue color to significantly distinguish them. For each pair of marmosets, the one dyed blue was annotated ID ‘1’ and the other one with normal white ears was annotated ID ‘2’, and each instance was also labeled 16 body locations. To train the 2D pose and identity predictor, we randomly selected 1354 frames as training data and the other 292 frames as test data.

### Marmoset3D

Ground truth of 3D poses are required in training denoising autoencoder and evaluating the accuracy of MarmoPose, which could be obtained by triangulating precisely hand-labeled 2D coordinates from multiple camera views at the same timepoint. To establish the Marmoset3D dataset, three human annotators labeled 522 3D ground truth, consisting of 140 instances triangulating from 560 images containing single marmoset and 382 instances triangulating from 191 images containing paired marmosets. Two annotators labeled the images first and then one annotator proofread the labels to ensure the accuracy.

### 2D pose tracking models training

MarmoPose employs a detection model adopted from Rtmdet(*36*) and a pose estimation model adopted from RTMPose(*37*) for 2D pose tracking. Both of the models were trained on the Marmoset3K dataset, with a division of 80% (2624 images, 3978 instances) for training and 20% (549 images, 841 instances) for test. The original images, with a resolution of (1080 *×* 1920 *×* 3), was downscaled to 640 *×* 640*×* 3 as the input of the detection model, then the bounding boxes predicted by detection model were cropped and resized to 512 *×* 512*×* 3 as the input of the pose estimation model. Training for both networks was conducted on a single NVIDIA RTX 4090 GPU using PyTorch 2.1.0.

### TensorRT deployment

After training the PyTorch models, we deployed them using TensorRT with the tools provided by MMDeploy to achieve faster inference speed. To accommodate inputs of various sizes, we configured dynamic input shapes, ranging from (1, 3, 640, 640) to (16, 3, 640, 640) for the detection model, and from (1, 3, 512, 512) to (64, 3, 512, 512) for the pose estimation model. FP16 precision was enabled to achieve real-time processing in the online module.

### Heuristic assignment of animal identities

Generally, the detection model will produce bounding boxes with duplicated labels. Given that each label instance appears only once in an image, we designed a heuristic assignment algorithm to refine the bounding boxes. In each iteration, we select the bounding box with the highest score from the remaining set, then we exclude any bounding boxes that have an intersection Over Union (IOU) overlap higher than 0.8 with the selected box. This process is repeated until the remaining set is empty or the number of selected bounding boxes matches the number of instances.

### SLEAP models training

For comparing with MarmoPose, we trained top-down models using SLEAP on the same Marmoset3K dataset. For the centroid model, we set ‘Max Stride’ to 32, ‘filters rate’ to 1.5 and ‘sigma’ to 2.5. Meanwhile, for the centered-instance model with id, we set ‘Max Stride’ to 64, ‘filters rate’ to 1.5 and ‘sigma’ to 2.5, the weights of loss for pose and identity are 999 and 0.01 respectively.

### Camera calibration

For camera calibration, we used the model and methods provided by Anipose(*24, 50*) to estimate the intrinsic and extrinsic parameters of multiple cameras. The calibration process involved placing a standard checkerboard pattern (11 *×* 9 squares with 45mm square size in this study) in the homecage and rotating it to various angles, then the recorded videos from different camera views are used for calibration.

In order to align the default coordinate system with actual spatial dimensions, we developed a labeling tool for users to define new coordinate system. This tool allows users to define a new coordinate system by marking three specific points in at least two camera views: the origin point, a point on the x-axis, and a point on the y-axis, then these points will be triangulated into 3D coordinates and cameras’ extrinsic parameters will be updated to fit the new coordinate system.

### Triangulation

In order to mitigate the impact of outlier 2D predictions, we first generated a set of candidate 3D poses through direct linear transformation (*24, 51*) across various combination of the camera views. Subsequently, a weighted reprojection error was computed for each candidate, and the one with the smallest error was selected as the final 3D pose. Where the weight of each point was determined by multiplying the likelihood of the point by the likelihood of the instance’s bounding box.

### Marmoset Skeleton Model

The marmoset skeleton model was established by manually measurement. Three normal adult marmosets were selected to measure the length of predefined body location pairs, and the median length was adopted as the reference value, as shown in Fig. 3c.

### Pose normalization

The reconstructed 3D poses are in the real spatial space where similar poses might have totally different coordinates, thus it is essential to normalize them into a unified coordinates system for effective DAE model training and subsequent analysis. We used three body locations to define the new coordinates system: the middle of spine (‘spinemid’) as the original point; the upper end of spine (’neck’) defining the x-axis; the lower end of spine (‘tailbase’) lying on the x-z plane. Then the original 3D poses are rotated into this unified coordinate system with standardize pose lengths by dividing each 3D coordinate with the distance between ‘spinemid’ and ‘neck’.

### Denoising autoencoder

Inspired by a pose estimation work on human(*39*), we adopted variational autoencoder to fill in the missing data caused by occlusion, which receives incomplete 3D poses as input and output the predicted complete 3D poses. Specifically, each pose is represented by a matrix *M_n×_*_3_, where *n* is the number of body locations (*n* = 16 by default), and the elements are the 3D coordinated of each body part in the real spatial space. We first normalized them into a unified coordinated system, and then flatten the matrix into a vector *V*_1*×*3*n*_ as the input of the DAE. With the Marmoset3D dataset, we generated incomplete 3D poses by randomly masking *i* (1 *≤ i ≤* 4) body locations to simulate the occlusion in real scenarios. These masked 3D poses and corresponding complete 3D poses are used as input and output of the DAE respectively during training.

In this study, the encoder of DAE is composed of 2 fully connected hidden layers, with 128 and 256 hidden units respectively. Symmetrically, the decoder also consists of 2 fully connected hidden layers with 256 and 128 hidden units. We used 40 latent dimensions to represent the input. The loss function consists of three parts: MSE calculating the difference between ground truth and reconstructed poses; *D_KL_* measuring the difference between the latent distribution and the standard normal distribution; and joint loss constraining the length between some body locations. Joint loss incorporates prior knowledge of the marmoset skeleton model into the model for guiding the reconstruction of missing body locations, making the DAE work better with few training data. The skeleton model is constructed by measuring the median length of distances between two body locations which are relatively fixed on adult marmosets, and each body part pair will be assigned with different weights based on their degrees of freedom. The DAE is trained with the Adam optimizer with a learning rate of 10*^−^*^3^.

### Real-time control module

The real-time control module in MarmoPose consists of two processes: (1) Prediction process. It reads the latest images from multiple live video streams cached by separate threads, then perform 2D detection and 3D triangulation and transmit the results to the main process. Customized predictors are used which stores the previous positions of marmosets for dynamic cropping. (2) Main process. It reads poses and images from the prediction process and display the real-time results, and also provides an interface for users to perform customized event detection and corresponding experimental control. An interface, receiving the latest 2D and 3D poses of marmosets, is provided in the real-time module, allowing users to perform customized event detection and corresponding experimental control. In the scenario of head orientation detection, we first got the head orientation of the marmoset by computing the vector from the middle of ‘left ear’ and ‘right ear’ to ‘head’ in the real space, then we defined the head orientation as ‘left’ if the angle between the head orientation vector and the normal vector of left side of the cage (i.e. x-z plane in the default coordinate system) is less than 45 degrees, and ‘forward’ if the angle is between 45 and 135 degrees, and ‘right’ if the angle is larger than 135 degrees.

Cameras produced live video streams at 25fps with 1920*×*1080*×*3 frame size, and frames are read by Real-Time Stream Protocol (RTSP). Latencies were evaluated on a computer with Intel Core i7- 13700K CPU, NVIDIA GeForce GTX 4090 GPU and 64GB Ram.

### Behavior map

For the creation of a behavioral map, a 32-minutes video clip containing 48000 poses of a single marmoset was analyzed. 97-dimensional features are selected for each 3D pose, including the non-locomoter movement (3×16=48 dims, derived by subtracting the ‘spinemid’ coordinated from each of the 3D pose without rotation), the locomotion speed (3×16=48 dims), and the hight of the ‘spinemid’ body location (1 dim). Then principal component analysis (PCA) was performed to reduce the features to 20 dimensions which account for 95.7% of the data variance. Subsequently, t-Distributed Stochastic Neighbor Embedding (t-SNE) was employed to generate the 2-dimensional embedding of all samples. By manually reviewing the poses at each local density peak, we identify 8 distinct postures of the marmoset, including climbing, jumping, squatting, proning, standing, walking, walking on sticks and hanging.

## Acknowledgment

This research was supported by Tsinghua University Non-human Primate Research Center (THU-NPRC). We thank Dr. Li Luo and veterinary and husbandry staff at THU-NPRC for help with this research.

## Author Contributions

C.C. and X.W. designed the study. C.C. developed the system. Z.H. helped develop the setup for video recording. C.C., R.Z., G.H, H.W. and L.T. labeled the dataset for model training and test. C.C. and X.W. wrote the manuscript.

## Data and materials availability

All data, code, and materials used in the analyses will be made publicly available on GitHub upon publication of this paper.

